# Effects of Gamma Irradiation Against Mating Competitiveness of Male *Culex quinquefasciatus* (Diptera:Culicidae) in Laboratory

**DOI:** 10.1101/727198

**Authors:** T Ramadhani, UK Hadi, S Soviana, Z Irawati, A Rahayu, Sunaryo, A Prastawa, A Mallongi

## Abstract

*Culex quinquefasciatus* is the main vector of lymphatic filariasis in Pekalongan City. Sterile Insect Tehnique (SIT) could be complementary vector control for filariasis. The key success of the technique depend on the ability of laboratory-reared sterile males with the wild-type females.The aim of the research was to determine the mating competitiveness, the fecundity and the fertility of sterile male Culex quinquefasciatus. The pupae of Cx. quinquefasciatus were gamma irradiated at the doses of 60Gy, 70Gy, and 80Gy, while unirradiated pupae were prepared as control. The mosquitoes emerging from the irradiated pupae could mate with a normal female in the cages. It were observed for the mean female laying eggs, the fecundity, the fertility and the mating competitiveness. The data were analyzed by one way ANOVA. The result showed that the irradiated *Cx. quinquefasciatus* at the doses tested did not affect on the fecundity and the mating competitiveness, but the fertility was disturbed (sterile). A dose of 70 Gy was the optimum dose or a fertility rate of 1.8% (98.2% sterile), and the value of competitiveness (C index) was 0.568. Based on the result, the irradiated *Cx. quinquefasciatus* can be recommended for semifield application.

## INTRODUCTION

Lymphatic filariasis is a communicable disease caused by filarial worms and transmitted through the bite of various kinds of mosquitoes. There are three species of worms causing lymphatic filariasis in Indonesia, namely: *Wuchereria bancrofti, Brugia malayi* and *Brugia timori* ^1^. The three species exist in Indonesia, but more than 70 percent of cases of filariasis in Indonesia were caused by *Brugia malayi* ^*2*^.

Pekalongan City is one of the endemic regions of lymphatic filariasis caused by *Wuchereria bancrofti* with *Culex quinquefasciatus* being the vector ^3^. Well-preserved places for *Culex quinquefasciatus* breeding, small coverage of Mass Drugs Administration (60.3 percent), people’s lack of awareness of environmental sanitation, the finding of filarial larvae in mosquitoes and vector’s resistence to insecticides are the factors that keeps filarial transmission going in Pekalongan City ^4,5,6^.

This also proves that the mass drug administration in the last five years has not effectively cut the chain of filarial transmission. An important effort that should be made is conducting surveilance of vector of filariasis as the basis for vector control in cutting the transmission chain. Vector control of filariasis in Indonesia has not been done in a dedicated way due to extremely complex issues. Fertile Insect Technique (SIT) is one of vector control efforts that can be made. SIT is a highly specific biological vector control technique which only affects target species. This technique only reduces the population size in the field, and not kills off the population, and graduated release may reduce the mosquito population ^7^. SIT pricinple is that male sterile insects (*Cx. quinquefasciatus*) are released in a considerable number to a target area. The released sterile male *Cx. quinquefasciatus* will compete with wild type males to mate wild type females. The mating between sterile male *Cx. quinquefasciatus* and wild type females is expected to produce no offspring, which eventually will reduce the population size of *Cx quinquefasciatus* in that area ^8^.

SIT is an environment-friendly, effective and potential vector control technique. This technique is also called species-specific control, by which vector is killed by the vector itself (autocidal technique)^9^. The technical procedure of this technique is relatively easy. Male mosquitoes are irradiated in the laboratory and then released to their habitat periodically. Irradiation may cause male mosquitoes to be sterile as it causes damages to spermatogenesis and stops the production of sperms (aspermia). Sperm inactivation also causes sterilization as sperms are not able to move to fertilize egg cell. The inability to mate serves as another sterilization factor as irradiation causes damages to somatic cells of internal genital organs and prevents egg cell fertilization^8^.

The application of sterile insect technique in Indonesia has been tried out on *Aedes aegypty* in vector control of high fever, and this technique has successfull reduced the size of mosquito population by up to 95.23 percent. This condition lasts for 3-6 months until high fever cases resurface. In the research on *Ae. aegypti* conducted by Batan, a gamma irradition dose of 70 Gy can be sterilized mosquitoes by up to 100 percent with a mating competitiveness value of 0.31, and a dose of 65 Gy can sterilize by 98.53 percent with a mating competitiveness value of 0.45^10^. On *Anopheles maculatus*, a dose of 110 Gy can sterilize by 97 percent with a mating competitiveness value of 0.65, and a dose of 120 Gy dose can sterilize by 99.99 percent ^11^. Setiyaningsih et al report that irradiation doses of 40 Gy, 50 Gy, 60 Gy and 70 Gy applied to *Cx quinquefasciatus* result in sterilization of 20.92 percent, 48.99 percent, 89.48 percent and 100 percent, respectively^12^.

SIT application for *Cx. quinquefasciatus* control has never been done in Indonesia, thus a preliminary test should be conducted for supporting the success of SIT application in the nature. Some preliminary tests are needed to support SIT application in the field, one of which is mating competitiveness of sterile male *Cx. quinquefasciatus*. This research was intended to evaluate the effect of gamma irradiation on fecundity, fertility and mating competitiveness of sterile male *Cx. quinquefasciatus* at the laboratory scale. It is expected that this research supports the feasibility of SIT application in lymphatic filarasis vector control.

## Materials and methods

### Ethics statement

The present study was performed with the approval of the Ethics in Research National Institute of Research and Development, Ministry of Health of Republic of Indonesia., who reviewed and authorized the procedures described in the study “Utilization of gamma irradiation on the sterility of the culex quinquefasciatus mosquito as an effort to control filariasis vector in Pekalongan City”. Mosquito rearing procedures and gamma irradiation were performed in the laboratory of entomology in center of Health Research and Development Banjarnegara Jl. Selamanik No 16 A and Center for Isotopes and Radiation Application, National Nuclear Energy Agency of Indonesia. Jl. Lebak Bulus Raya 49, Jakarta 12070, Indonesia. Mosquito eggs were collected from Pekalongan city central Java.

### Sample Mosquitoes

The research sample consisted of *Culex quinquefasciatus* pupae of second to third generation from the field isolate in Pabean Subdistrict, Pekalongan City kept at the Laboratory of Entomology of the Center for Research and Development of Animal-Borne Disease Control of Banjarnegara. The colony was from the *Culex spp* larvae in Pabean Subdistrict, Pekalongan City, and then kept at the laboratory, and the larvae were fed with uncrushed dog feed (Pedigree®). The administration of larva feed was adjusted to the age, after instar 3 was moved to cultivation tray sized 27 cm × 35 cm × 5 cm with a density of 400-600 mosquitoes/tray. The larvae were raised until they grew into pupae. The pupae raised every day were taken and put into mosquito cage. The emerging mosquitoes in the cage were identified and separated by *Cx. quinquefasciatus* and given 10% sugar solution and marmot blood to enable them to lay eggs. The eggs produced were incubated and raised until they turned into pupae. The male and female pupae derived from the colonization at the laboratory were separated (male pupae are smaller than female pupae) using a sieving device (90-95 percent small pupae are males) according to the method of Nurhayati et al (2010) ^13^. The identified male pupae were inserted into a bottle, kept in a cool box and carried to PAIR BATAN Jakarta to be irradiated. Female pupae were inserted into a 40 cm × 40 cm × 40 cm sized mosquito cage and then kept at the Laboratory of Entomology of the Center for Research and Development of Animal-Borne Disease Control of Banjarnegara before test.

### Gamma irradiation

Irradiation was conducted at the Center for Technology of Isotope and Radiation BATAN, Jakarta. Irradiation was carried out when *Cx. qinquefasciatus* pupae were ≥15 hours old using Gamma Cell-220 irradiator and based on the the position of radioactive sources around the material, so more uniform absorption doses were obtained. The sample were placed in plastic pots with lower diameter of 4.5 cm, upper diameter of 6.5 cm and height of 6.5 cm, filled with 20 ml of water. Every plastic pot was filled with 200 pupae to which irradiation doses of 60, 70 and 80 Gy were administered separately and alternately. The doses were determined based on the results of dose test of previous research. The control group consisted of non-irradiated male *Cx. quinquefasciatus* having been prepared at the Laboratory of Entomology of the Center for Research and Development of Animal-Borne Disease Control of Banjarnegara at the same age of that of the irradiated pupae. After the gamma irradiation process, male *Cx. quinquefasciatus* pupae were raised until they became mature, and observation was conducted on their fertility, fecundity and mating competitiveness.

### Fecundity and fertility

Mature male *Cx. quinquefasciatus* (n=25) on which gamma irradiation was administered were mated with normal female mosquitoes (n=25) in a cubic cage sized 30 cm × 30 cm × 30 cm. The control group consisted of normal male *Cx. quinquefasciatus* (n=25) that were mated with normal female mosquitoes (n=25). After three days of mating period, the male mosquitoes were released from the cage using aspirator. Female mosquitoes were fed with blood (marmot) until they were full, and then individually moved to a transparent plastic vial tubbe for oviposition process^14^. This treatment was repeated four times. The observation of the number of eggs, eggs that hatched and eggs that did not hatch was conducted manually using stereo microscope with a 4x magnification. Eggs with open operculum were eggs that hatched, while eggs that did not hatch were marked with closed operculum^15^.

### Mating competitiveness

The observation of mating competitiveness was conducted at the Laboratory of Entomology of the Center for Research and Development of Animal-Borne Disease Control of Banjarnegara at a temperature of 22-25 °C and humidity of 70-79 percent. After the irradiation process, each mosquito was inserted into the cage by taking into account the ratio between irradiated males, normal males and normal females. As many as 25 male *Cx. quinquefasciatus* and 25 female *Cx. quinquefasciatus* were used in every mating combination with four repetations. The mating process took place naturally, where males and females were placed in a 30 cm × 30 cm × 30 cm sized cubic cage for 2-3 days for maximizing the mating. The irradiated male mosquitoes were marked with code R, while the normal female and male mosquitoes were marked with code N. Then, all mosquitoes were mated following the combinations in the method of Bellini et al ^16^ as shown in Table 1.

**Table 1.**
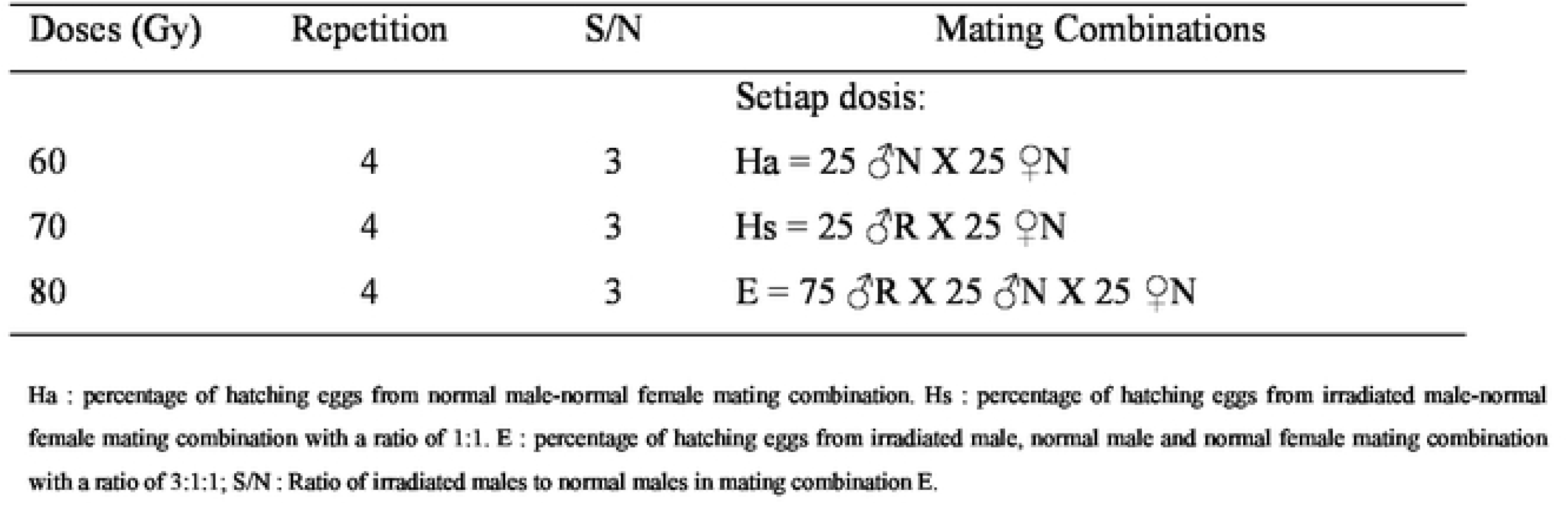
Mating combinations of *Cx. quinquefasciatus* for the observation of mating competitiveness

The mosquitoes in the mating cage were given 10% sugar solution. For five days starting from the fourth day— the mosquitoes were given marmot blood for one hour each day. Gravid female mosquitoes were taken individually and inserted into 150 ml sized paper glass filled with 50 ml of 10% straw water as a place to lay eggs. The observation of the number of eggs and hatching eggs was conducted every day for five days. The observation was conducted for one gonotrophic cycle.

### Data analysis

The parameters measured in this research were the number of gravid female mosquitoes, number of eggs produced by female mosquitoes (fecundity), number of hatching eggs (fertility) and the value of mating competitiveness obtained from Fried index (C index). Fertility rate was calculated by comparing the number of hatching eggs and the number of all eggs. The mating competitiveness value or C index was obtained from the equation according to the method of Bellini et al (2013).

The data were analyzed using SPSS version 16, and one way-analysis of variance (ANOVA) was used for evaluating the effect of the gamma irradiation doses on fecundity, fertility and mating competitiveness with p value of <0.05. Duncan’s post hoc test was used for further analysis of the comparison between means.

## RESULTS

### Effects of irradiation on fecundity

The data of fecundity and number of normal female *Cx. quinquefasciatis* mosquitoes successfully laying eggs by mating with irradiated males and normal males (non-irradiated) as control group are presented in Table 2. The percentage of female mosquitoes successfully laying eggs was 40 percent in the control group and 39-43 percent in the treatment group, and there was no difference statistically (F = 2.201; *p* = 0,099).

**Table 2.** Fecundity of female *Culex quinquefasciatus* in mating with irradiated male mosquitoes with different dosages

| Doses (Gy) | Average % of female mosquitoes laying eggs | Number of eggs | Average number of eggs per mosquito ± SD (95 % CI) |
| --- | --- | --- | --- |
| Control | 40.00<sup>a</sup> | 4528<sup>a</sup> | 111.8±10.2 (95.5-128.1)<sup>a</sup> |
| 60 | 39.00<sup>a</sup> | 4264<sup>a</sup> | 109.3±26.5 (67.5-151.4)<sup>a</sup> |
| 70 | 43.00<sup>a</sup> | 5160<sup>a</sup> | 120.7±11.1 (103.1-38.4)<sup>a</sup> |
| 80 | 41.00<sup>a</sup> | 3939<sup>a</sup> | 96.4±27.5 (52.7-140.2)<sup>a</sup> |

The different superscripts in the same columns show significant difference ( $p < 0.05$ ) in ANOVA Duncan's post hoc test.

### Effects of irradiation on Fertility

Fertility rate is the percentage of the number of hatching eggs out of the number of eggs produced by female mosquitoes mating with sterile or normal male mosquitoes. The fertility rates of female mosquitoes mating with irradiated and control male mosquitoes are shown in Table 3. Normal female mosquitoes mating with normal males had the highest fertility at 94.1 percent, while those mating with irradiated males had fertility of around 1.3-4.8 percent. The higher the irradiation dose, the lower the fertility rate. The results of the statistical test show a significant difference in the fertility rate of treatment group and control group (F = 896.614; *p* = 0.00).

**Table 3.** Fertility rate of female *Culex quinquefasciatus* mating with irradiated male mosquitoes at different doses

| Doses (Gy) | Average number of eggs | Average number of hatching eggs | Average % of hatching eggs ±SD (95 % CI) |
| --- | --- | --- | --- |
| Control | 1132.00 | 1076.25 | 94.1±5.4 (85.6-102.7)<sup>b</sup> |
| 60 | 1066.00 | 51.17 | 4.8±1.5 (2.4-7.3)<sup>a</sup> |
| 70 | 1290.00 | 23.22 | 1.8±0.6 (1.0-2.6)<sup>a</sup> |
| 80 | 984.75 | 12.80 | 1.3±0.6 (0.4-2.4)<sup>a</sup> |

The different superscripts in the same columns show significant difference ( $p < 0.05$ ) in the ANOVA Duncan's post hoc ANOVA test

### Effects of irradiation on mating competitiveness

The mating competitiveness values of sterile *Cx. quinquefasciatus* at various irradiation doses are presented in Table 4. The highest C index value was obtained at the irradiation dose of 70 Gy (C index = 0.53), while the lowest was at the irradiation dose of 80 Gy (C index = 0.36). The statistical test results show no significant difference in the mating competitiveness of sterile male *Cx. quinquefasciatus* at different irradiation doses (F = 0.526; p > 0.05).

**Table 4.** The Mating Competitiveness Values of sterile *Cx. quinquefasciatus* at different irradiation doses with mating combinations at the laboratory scale

| Doses (Gy) | Percentage of the hatching of mating combination ♂R: ♂N: ♀N |  |  | S/N | C index |
| --- | --- | --- | --- | --- | --- |
|  | 0 : 25 : 25 | 25 : 0 : 25 | 75 : 25 : 25 |  |  |
| 60 | 91.52 | 3.15 | 43.65 | 3 | 0.44<sup>a</sup> |
| 70 | 91.54 | 2.27 | 38.32 | 3 | 0.53<sup>a</sup> |
| 80 | 91.51 | 1.05 | 45.13 | 3 | 0.36<sup>a</sup> |

The different superscripts in the same columns show significant difference ( $p < 0.05$ ) in the ANOVA Duncan's post hoc ANOVA test

## DISCUSSION

The ability of mosquitoes to lay eggs is related to the structures and functions of the reproductive organs. Mosquitoes with healthy condition of reproductive organs and ability to function normally tend to have more successful oviposition than the otherwise. Mosquitoes may be unable to lay eggs because of their ovaries’ inability to produce eggs or to have oviposition. The two factors are related to the biological aspect, especially the factor that is related to abnormalities of the organ function due to genetic factors^17^.

Table 2 shows the number of eggs produced by female *Cx. quinquefasciatus* mating with irradiated males. The highest number was produced at an irradiation dose of 70 Gy (5,160 eggs), and the lowest number at 80 Gy (3,939 eggs). The results of the statistical analysis show that there was no significant difference between the number of eggs produced by female *Cx. quinquefasciatus* at various irradiation doses and the control group (F = 1,544; *p* = 0.254; p > 0.05).

The largest number of eggs produced by female *Cx. quinquefasciatus* per oviposition was at the irradiation dose of 70 Gy (120.7 ± 11.1) and the lowest was at the dose of 80 Gy (96.4 ± 27.5). The results of the observation show that there was no difference between the average number of eggs per female *Cx. quinquefasciatus* at different irradiation doses and the control group (F = 0.907; *p* = 0.498; p > 0.05) (Table 2). These results are in line with the results of the research by Sasmita and Ernawan ^18^ on *Ae. aegypti* and *An. maculatus*^19^.

The number of eggs produced by female mosquitoes is affected by the amount of blood sucked, biological factor and ecological factor^17^. According to Clements, to produce 85.5 eggs, on average, a mosquito needs 3-5 mg of blood. Eggs will not be produced if the amount of blood sucked is less than 0.5 mg ^20^. The size of the mosquito’s body is a biological factor, which affects the size of its abdomen and ovary. Larger ovary, which functions as a place in which eggs are produced, has bigger capacity and productivity. Large mosquitoes tend to have larger abdomens. In addition, the bigger the amount of blood is sucked, the bigger the number of the eggs is produced^21^.

The eggs produced by female *Cx. quinquefasciatus* show that the insemination process went on, but the embryo formed could not survive due to the dominant lethal mutation properties carried by irradiated male mosquito sperm cells. Dominant lethal mutation does not inhibit the process of male nor female gametogenesis, and there is no inhibition in the zygot formation either, but the embryo will experience death^9^. The dominant lethal mutation can be seen from the decreasing fertility rate with the increase in the gamma irradiation doses administered ^22^.

The control group had high fertility rate because the sperm cells transfered by male mosquitoes to female spermatheca in the mating process were normal. The meeting of normal sperm cell and egg cell of a female mosquito will produce a fertile egg ^20^.

Setiyaningsih et al reports that SIT application has an effect on the decrease in the fertility of the eggs of *Ae. aegypti*, both in and out of home ^20^. The decrease in the percentage of fertility is due to the mating between sterile male *Cx. quinquefasciatus* and normal females in the nature. Sterile male mosquitoes will transfer sterile sperm cells to the spermatheca of female mosquitoes and produce sterile eggs. Sterile eggs may also be formed when sterile male mosquitoes fail to mate with female mosquitoes due to the morphological changes in the male mosquitoes’ sex organs because of irradiation process. The morphological changes in the sex organs of male mosquitoes may hinder the transfer of sperm cells to female mosquitoes and prevent egg cell fertilization^20, 23^.

Pupal stadium is a developmental stadium where young organs transform or develop into mature organs^24^. In this stadium, spermatogenesis normally takes place. A little dose of radiation (65-70 Gy) may have caused sterilization already. In the permatogenesis process, the sperm cells split rapidly. If the irradiation administered on the sperm cells, some changes will occur and abnormal sperm cells are produced. Abnormal sperm cells have small heads, short tails and low mobility, while normal sperm cells are bigger and have higher mobility ^23^.

The percentage of egg hatching of female mosquitoes used in this research was fairly good. It reached 94.1 percent. This is different from the research by Sasmita and Ernawan on *Aedes aegypti*^18^, where the percentage of egg hatching was 50.6 percent. The variation in the gamma irradiation doses affects the fertility and sterilization of *Cx. quiquefasciatus*, which is in line with Setiyaningsih et al ^12^. Shetty et al report the effect of gamma irradiation dosage of 0-50 Gy on fertility, egg hatching, occurance and age of *Ae. Aegypti* ^27^.

Some eggs do not hatch because there is no fertilization between sterile sperm cells and egg cells because male *Cx. quiquefasciatus* are unable to perfectly copulate with normal females as their genital organs change, preventing the sperm cells to be delivered perfectly (Helinski *et al* 2008). Decreased egg fertility also shows the ability of sterile male *Cx. quiquefasciatus* in competing with normal male *Cx. quiquefasciatus* in the nature in finding mates for mating.

High mating competitiveness shows that the gamma irradiation dose of Co-60 administered in the pupal stadium did not have any effect on the mating competitiveness. Every species needs a certain optimum gamma irradiation dose for the sterilization of eggs without affecting its mating competitiveness ^22,26^

In this research the irradiation dose did not have any effect on the mating competitiveness statistically as it was possibly caused by relatively small range of the doses tested. Some research has proven that higher doses will cause negative effect on mating competitiveness^26^. This pattern is also found in *Ae. Aegypti*^*15*^ and *An arabiensis* ^*23*^. Higher irradiation doses (> 120 Gy) needed for sterilizing males may also cause such males to be unable to transfer sperm cells into female mosquitoes ^13^.

Sterile males’ success in mating may be caused by their ability to compete with normal males in getting mates. The main factor of the decrease in mating competitivenss in SIT is gamma irradiation sterilization process. Ionizing gamma irradiation may cause damages to somatic cells, which may also deteriorate male *Ae. aegypti*’s form ^16,25^. Low physical form may also worsen males’ ability to mate with females. The results of some previous research show that the mating competitiveness of male mosquitoes has a negative correlation with the increase in gamma irradiation dose ^16,18^.

A dose of 70 Gy is an optimum dose to be used at a fertility rate of 1.8 percent, with competitiveness value of 0.53. This shows that the irradiation dose of 70 Gy can decrease the egg hatching rate by 98.2 percent. Mating competitiveness value or C Index is useful for determining the number of sterile male mosquitoes spread in the nature following the method of Sasmita and Ernawan^18^. In this research the mating competitiveness of *Cx. quinquefasciatus* was 0.53. This means that the number of sterile males released should be at least twice of the population of males in the nature in order to increase mating possibility with normal females. Our results that wild *Cx. quinquefasciatus* males irradiated at the optimum dose (70 Gy) can compete successfully with unirradiated male for wild females in laboratory.

## CONCLUSION

Irradiation in *Culex quinquefasciatus* pupae at a dose of 60-80 Gy affects fertility, but does not affect the fecundity and mating competitiveness. A dose of 70 Gy is an optimum dosage as it may yield fertility rate of 1.8 percent (98.2 percent sterile) and mating competitiveness value of 0.53 (i.e. that the number of sterile male mosquitoes released should be twice the number of male mosquitoes in the population).

The results of this research can be used as the baseline for further testing at semi-field and limited-field scale and for assessing SIT feasibility in the vector control of lymphatic filariasis in Pekalongan City.

## ACKNOWLEDGMENT

Authors thank for head of Research and Development of Banjarnegara-Sourced Disease Control Health. This study was supported by Health Research and Development Agency. We also thank support from the entomology Laboratory Research and Development of Banjarnegara and PAIR BATAN Jakarta as well as for those who helped during the author conduct his research in the laboratory observation.

## Authors’ contributions

TR, UKH, SS and ZI carried out the experiments. AR and S contributed to the rearing, sexing and irradiating. TR, UKH drafted the initial manuscript or performed the statistical analysis and reviewed the early version. UKH, SS and ZI helped to draft the last version of the manuscript. TR designed the experiment and supervised the entire work. All authors read and approved the final manuscript.

## Data Availability

All relevant data are within the paper and its supporting information files

## REFERENCES

1. [Pusdatin] Pusat Data dan Informasi Kementerian Kesehatan RI. Situasi Filariasis di Indonesia Tahun 2015., Infodatin ISSN 2442-7659. 2016

2. [Ditjen PP&PL]. [Ditjen PP&PL]. Direktorat Jenderal Pengendalian Penyakit dan Penyehatan Lingkungan. Laporan Bulanan dan tahunan Subdit filariasis Ditjen PP& PL Jakarta 2009

3. Ramadhani R, Suyoko, Sumarni S. *Culex quinquefasciatus* sebagai Vektor Utama Filariasis Limfatik yang disebabkan *Wuchereria bancrofti* di Kelurahan Pabean Kota Pekalongan., Jurnal Ekologi Kesehatan., 9(3):1303–1310, 2010

4. Windiastuti IA, Suhartono, Nurjazuli. Hubungan Kondisi Lingkungan Rumah, Sosial Ekonomi, dan Perilaku Masyarakat dengan Kejadian Filariasis di Kecamatan Pekalongan Selatan Kota Pekalongan. Jurnal Kesehatan Lingkungan Indonesia., 12(1):51–57, 2013

5. Ramadhani T, Wahyudi BF. Keanekaragaman dan Dominasi Nyamuk di Daerah Endemis Filariasis Limfatik Kota Pekalongan. Jurnal Vektor Penyakit., 9(1):1–8, 2015

6. Wahyudi FW, Pramestuti N. Kondisi Filariasis Pasca Pengobatan Massal di Kelurahan Pabean Kecamatan Pekalongan Utara Kota Pekalongan. Jurnal Balaba., 12(1):55–60, 2016

7. Dyck VA, Hendrichs J, Robinson AS. Sterile Insect Technique Principles and Practice in Area-Wide Integrated Pest Management., The Netherlands: Springer

8. Knipling EF. 1955. Possibilities of Insect Control or Eradication Through the Use of Sexuality Sterile. J.Econ. Entomol., 48: 459–462, 2005

9. Sutrisno S. Dasar Penerapan Teknik Serangga Mandul untuk Pengendalian Hama pada Kawasan yang Luas., Jurnal Ilmiah Aplikasi Isotop dan Iradiasi., 2(2):35–47, 2006

10. Nurhayati S, Santoso B, Rahayu A. Pengendalian Populasi Nyamuk *Aedes aegypti* dan *Anopheles sp* sebagai Vektor Demam Berdarah Dengue (DBD) dan Malaria dengan Teknik Serangga Mandul (TSM). Seminar Nasional Keselamatan Kesehatan dan Lingkungan VI Jakarta., 15–16 Juni 2010 PTKMR-BATAN, FKM-UI, KEMENKES-RI, 2010

11. Widiarti. Pengembangan Teknik Serangga Mandul dengan Iradiasi Gamma dalam Upaya Pengendalian Vektor Malaria Di Kabupaten Kebumen Jawa Tengah. Laporan akhir Penelitian Ristek (unpublish) 2010

12. Setiyaningsih R, Widiarti, Heriyanto H. Pengaruh Pelepasan nyamuk Jantan Mandul terhadap Fertilitas dan Perubahan Morfologi Telur *Aedes aegypti*. Jurnal Vektora., 7(2):71–78, 2015

13. Nurhayati S, Santoso B, Rahayu A, Devita T. Pengaruh Radiasi sinar Gamma Terhadap Daya Saing Kawin Nyamuk Aedes aegypti sebagai Vektor Demam Bedarah Dengue (DBD)., Prosiding Seminar Nasional Keselamatan dan Lingkungan V, Depok. 2009

14. Zhang D, Lees RS, Xi Z, et al. Combining the Sterile Insect Technique with the Incompatible Insect Technique: III-Robust Mating Competitiveness of Irradiated Triple Wolbachia-Infected Aedes albopictus Males under Semi-Field Conditions. PloS ONE., 10(4):1–13, 2016

15. Zheng ML, Zhang DJ, Damiens DD, et al. Standard operating procedures for standardized mass rearing of the dengue and chikungunya vectors *Aedes aegypti* and *Aedes albopictus* (Diptera: Culicidae)-II-Egg storage and hatching. Parasites & Vectors., 10:1–7, 2015

16. Bellini R, Balestrino F, Medici A, et al. Mating Competitiveness of *Aedes albopictus* Radio-Sterilized Males in Large Enclosures Exposed to Natural Conditions. Journal Med Entomol., 50(1):94–102, 2013

17. Iswanto, Mardihusodo SJ, Baskoro T. Tabel Kehidupan dan Fekunditas *Culex quinquefasciatus* Say (Diptera: Culicidae) Kota Yogyakarta dan Semarang di Laboratorium. Jurnal Sain Kesehatan., 17:89–104, 2004

18. Sasmita HI, Ernawan B. Kualitas Nyamuk Jantan Mandul Aedes aegypti L. Hasil Iradiasi Gamma: Efek Iradiasi Pada Fase Pupa dan Dewasa. Jurnal Ilmiah Aplikasi Isotop dan Radiasi., 10(2): 149–158, 2014

19. Clements AN. The Physiology of Mosquitoes. New York : A Pergamon Press Book. 393p 1963

20. Setiyaningsih R, Agustini M, Rahayu A. Pengaruh Pelepasan nyamuk Jantan Mandul terhadap Fertilitas dan Perubahan Morfologi Telur *Aedes aegypti*. Jurnal Vektora., 7(2):71–78, 2015

21. Herms BJ. Medical Entomology 4^th^ ed. New York : The Mc Millan Co 1950

22. Nurhayati S, Yunianto B, Ramadhani T, et al. Controlling *Aedes aegypti* Population as DHF Vector with Radiation Based-Sterile Insect Technique in Banjarnegara Regency, Central Java. Jurnal Sains dan Teknologi Nuklir Indonesia., 14(1): 1–10, 2013

23. Helinski MEH, Parker AG, Knols BGJ. Sperm quantity and size polymorphism in un-irradiate male of the malaria mosquito *Anopheles arabiensis* Patton. Acta Tropica., 109: 64–69, 2008

24. Hoper GHS. Competitiveness of Gamma Sterilized Males of the Mediteranean Fruit Fly : Effect of Irradiating Pupae or Adult Stage and of Irradiating Pupae in Nitrogen. J.Econ.Entomol., 64:464–468 in Area-Wide Integrated Pest Management eBook 1976

25. Calkins CO and Parker A G. Sterile insect quality In Dyck VA and Robinson AS (Eds.) Sterile Insect Technique Principles and Practice in Area-Wide Integrated Pest Management. The Netherlands: Springer p 269–296, 2005

26. Helinski MEH, Knols BGJ. Mating competitiveness of male *Anopheles arabiensis* mosquitoes irradiated with a partially or fully sterilizing dose in small and large laboratory cages. J.Med.Entomol., 45(1):698–705, 2008

27. Shetty V, Shetty NJ, Harini BP, et al. Effect of gamma radiation on life history traits of *Aedes aegypti* (L.). Parasite Epidemiology and Control., 1(2): 26–35, 2016

